# Cohesin chromatin loop formation by an extrinsic motor

**DOI:** 10.1101/2023.11.30.569410

**Authors:** Thomas M. Guérin, Christopher Barrington, Georgii Pobegalov, Maxim I. Molodtsov, Frank Uhlmann

## Abstract

The ring-shaped cohesin complex topologically entraps two DNAs to establish sister chromatid cohesion^1–3^. Cohesin also shapes the interphase chromatin landscape with wide-ranging implications for gene regulation^4–7^, which cohesin is thought to achieve by actively extruding DNA loops without topologically entrapping DNA^8–11^. The ‘loop extrusion’ hypothesis finds motivation from *in vitro* observations^12–14^ – whether this process underlies *in vivo* chromatin loop formation remains untested. Here, using the budding yeast *S. cerevisiae*, we generate cohesin variants that have lost their ability to extrude DNA loops but retain their ability to topologically entrap DNA. Analysis of these variants suggests that *in vivo* chromatin loops form independently of loop extrusion. Instead, we find that transcription promotes loop formation, as well as acts as an extrinsic motor that expands these loops and defines their ultimate positions. Our results necessitate a re-evaluation of the loop extrusion model and point to an alternative mechanism for cohesin-dependent chromatin organisation. We propose that cohesin, akin to sister chromatid cohesion establishment at replication forks, forms chromatin loops by DNA-DNA capture at places of transcription, thus unifying cohesin’s two roles in chromosome segregation and interphase genome organisation.

## Introduction

Cohesin was first identified for its role in sister chromatid cohesion, holding DNA replication products together to allow their faithful distribution to daughter cells during cell divisions^15–17^. The ring-shaped cohesin complex achieves this by topologically entrapping both sister chromatids^1–3^. At a molecular level, cohesin sequentially and topologically entraps two DNAs, with preference for double stranded DNA (dsDNA) followed by single stranded DNA (ssDNA). This configuration matches that at DNA replication forks where the dsDNA leading strand product lies juxtaposed to the unwound ssDNA lagging strand. Second ssDNA capture is labile but turns stable by ssDNA to dsDNA conversion during lagging strand DNA synthesis and concomitant cohesin acetylation^2,18^.

In addition to sister chromatid cohesion, cohesin plays a key role in interphase genome organisation. Cohesin establishes chromatin loops, as well as demarcates topologically associating domains (TADs) within which chromatin interactions are enriched^4–7^. Cohesin-dependent loops are often seen between chromosomal CCCTC-binding factor (CTCF) binding sites, where cohesin accumulates in mammalian genomes. Acutely removing cohesin from interphase nuclei causes only limited transcriptional changes^6,7,19,20^. At the same time, changing cohesin-dependent chromatin interactions by altering CTCF binding sites profoundly impacts gene regulation in the long term^21–26^.

In principle, cohesin could form interphase chromatin loops by sequential topological capture of DNA sequences along the same chromatin chain. However, over recent years, an alternative model for loop formation by cohesin has gained prominence, loop extrusion^8–11^. According to the loop extrusion hypothesis, cohesin generates a small DNA loop that cohesin then enlarges using intrinsic DNA motor activity. Single molecule *in vitro* experiments have strikingly illustrated cohesin-mediated DNA loop extrusion^12–14^. However, very low external forces stall loop extrusion, and it remains uncertain how cohesin might navigate the complex *in vivo* chromatin landscape. DNA-bound obstacles are portrayed either as surmountable^27^ or as barriers^28,29^. Whether *in vivo* chromatin loops and TADs indeed form by loop extrusion has not yet been experimentally tested.

## Loop extrusion by budding yeast cohesin

We purified recombinant budding yeast cohesin, as well as its loader, following overexpression in budding yeast (Fig. 1a and Supplementary Fig. 1a)^30^. When added to 48.5 kb long λ-DNA, loosely tethered to a flow cell surface, and stained with SYTOX Orange, we observed efficient DNA loop extrusion (Fig. 1b). The observed loop extrusion rate (620 ± 440 bp s^-1^, mean ± s.d.) was only slightly lower than previously reported rates of human and fission yeast cohesin (Supplementary Fig. 1b)^12–14^. Consistent with previous observations, loop extrusion by budding yeast cohesin strictly depended on the presence of the cohesin loader (Fig. 1b). Thus, the ability to perform *in vitro* loop extrusion extends to budding yeast cohesin.

**Fig. 1.**
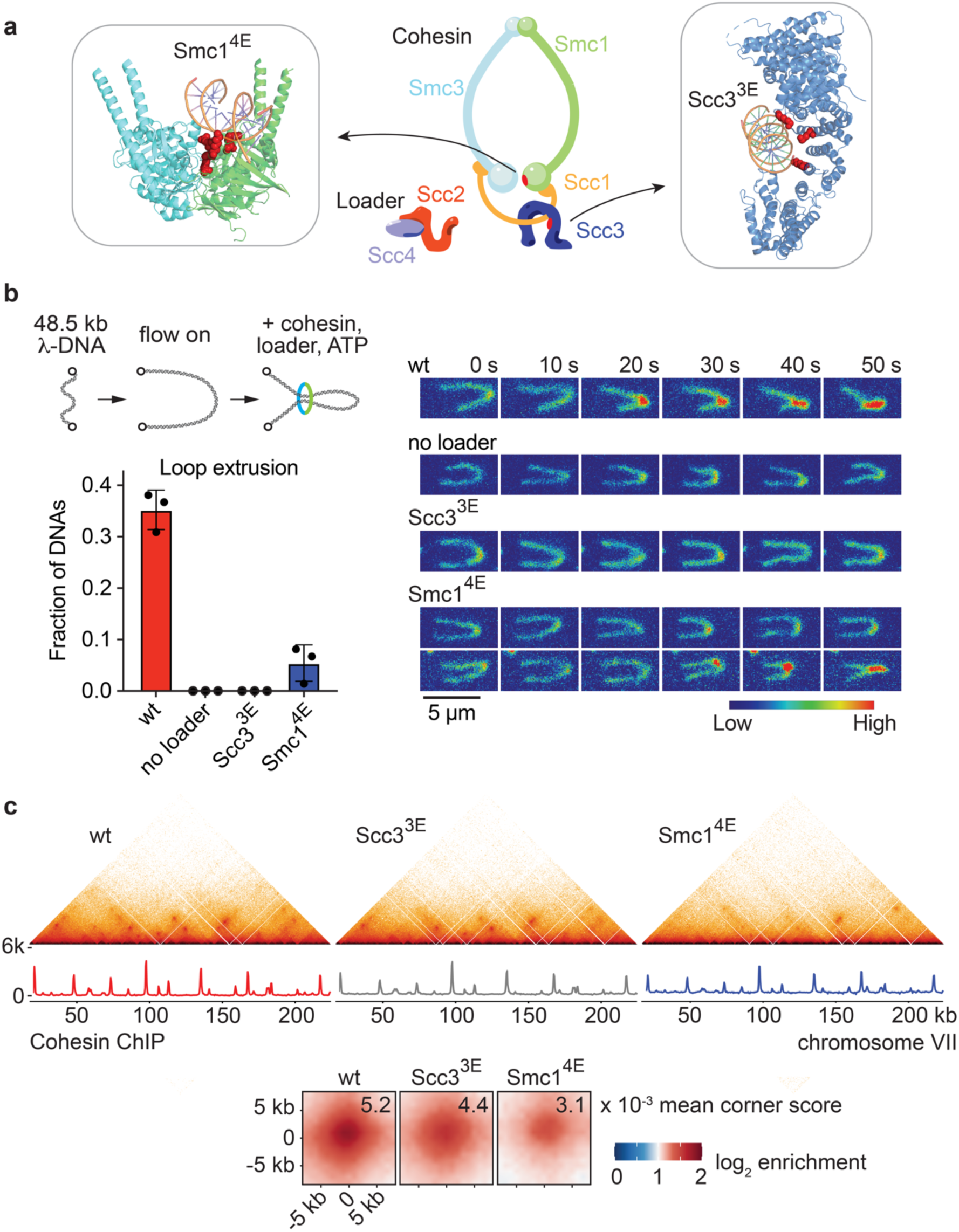
Chromatin loop formation by loop extrusion deficient cohesin. **a,** Overview schematic of cohesin and its loader, and structures of the Scc3 (PDB: 6H8Q)^36^ and Smc1 (PDB: 6ZZ6)^79^ subunits bound to DNA, highlighting the amino acids that were mutated to glutamates to generate Scc3^3E^ (K423E, K520E, K669E) and Smc1^4E^ (R53E, R58E, N60E, K63E). **b,** Schematic representation of an *in vitro* loop extrusion assay. Loop extrusion efficiencies of wild type (wt) cohesin in the presence and absence of loader, as well as of Scc3^3E^- and Smc1^4E^- cohesin, were measured in three independent repeat experiments. Individual data points are presented, bars indicate the mean and error bars the standard deviation (n_wt_ = 201, n_no loader_ = 224, n_Scc3_^3E^ = 301, n_Smc1_^4E^ = 260). Example time lapse recordings of loop extrusion. The DNA is stained with SYTOX Orange and shown using an arbitrary linear intensity scale. Scale bar, 5 μM. **c,** 500 bp-resolution merged micro-C contact maps, as well as corresponding calibrated cohesin ChIP-seq traces, from two independent experiments of G2/M arrested cells harbouring wt or loop extrusion deficient cohesin. Cohesin ChIP used Smc3-Pk_3_ in the wt and Scc3^3E^ strains, or Smc1^4E^-Pk_3_ with a wt Smc1-Pk_3_ strain included for normalization. Aggregate chromatin structure is shown in wt, Scc3^3E^ and Smc1^4E^ strains at loops detected by chromosight^76^ and linked to cohesin anchors in the wt strain (n = 1060). Mean corner scores are indicated.

Several molecular models have been proposed to explain loop extrusion^14,31–34^. While the actual mechanism remains to be ascertained, DNA binding by cohesin’s Scc3 subunit plays a central role in all proposals. Consistent with such a contribution, charge reversal mutations in human Scc3^STAG1^ render cohesin unable to extrude DNA loops, as did mutations to a DNA binding surface on the Smc1 ATPase head^33,35^. We replaced three budding yeast Scc3 lysine residues known to contact DNA with glutamate (Scc3^3E^)^36^, as well as made four glutamate exchanges on Smc1 (Smc1^4E^; Fig. 1a). These alterations did not noticeably change cohesin’s overall affinity to DNA, as measured in a DNA electrophoretic mobility shift assay. Scc3^3E^- and Smc1^4E^-cohesin also retained their ability to entrap DNA in an ATP-dependent, salt-resistant manner, albeit with reduced efficiency, especially in the case of Smc1^4E^-cohesin (Supplementary Fig. 1c,d)^30^. Monitoring loop extrusion in real time, while DNA molecules were stretched by mild liquid flow, revealed that Scc3^3E^-cohesin had completely lost its ability to extrude DNA loops, while Smc1^4E^-cohesin showed a greatly reduced loop formation frequency (Fig. 1b). To assess loop extrusion in a more sensitive assay, we incubated cohesin and its loader with DNA in the absence of flow, then applied flow solely to visualise the resultant DNA loops. In this assay, Smc1^4E^-cohesin formed DNA loops with close to half the wild type efficiency, while Scc3^3E^-cohesin remained loop extrusion deficient (Supplementary Fig. 1e).

## Life without loop extrusion

We next generated budding yeast strains expressing Scc3^3E^ or Smc1^4E^ as the sole source of these respective cohesin subunits. The resultant strains displayed no noticeable growth defects nor sensitivities to genome damaging agents (Supplementary Fig. 2a). When arrested at G2/M (by release from α-factor synchronisation into nocodazole-containing medium) Scc3^3E^ and Smc1^4E^ cells displayed slight sister chromatid cohesion defects, compared to a wild type control (Supplementary Fig. 2b), perhaps because of the compromised ability of the mutant cohesin complexes to load onto DNA^36^. This interpretation found support when we measured *in vivo* cohesin loading using calibrated ChIP-sequencing. Scc3^3E^- and Smc1^4E^-cohesin associated with chromosomes in a pattern indistinguishable from wild type cohesin, but at reduced levels (Fig. 1c and Supplementary Fig. 2c).

Budding yeast display a prominent cohesin-dependent looping pattern between neighbouring cohesin binding sites that has been visualised by micro-C^37^. We observed these chromatin loops similarly in a wild type strain, as well as in strains harbouring loop extrusion deficient Scc3^3E^- or Smc1^4E^-cohesin (Fig. 1c). Quantification of the loop signals (corner scores) showed a reduced loop intensity in the Scc3^3E^- and Smc1^4E^-expressing strains, as might be expected from reduced cohesin levels, but loops remained clearly discernible. Loops were weaker in Smc1^4E^- than in Scc3^3E^-cells, meaning that loop strength correlated with cohesin’s *in vitro* ability to entrap DNA more than with its ability to perform loop extrusion (Supplementary Fig. 2d). We conclude that cohesin complexes that cannot extrude DNA loops, especially Scc3^3E^-cohesin, nonetheless support yeast cell growth and cohesin-linked chromatin loop formation. These findings imply that cohesin forms chromatin loops by a mechanism different from loop extrusion.

## Transcription promotes chromatin loop formation

Transcription has been implicated in establishing chromatin architectural patterns^38–40^, and it is thought that RNA polymerases do so by forming barriers to loop extruding cohesins^40,41^. To revisit this phenomenon, we designed a yeast strain in which both the Rpb1 and Rpb3 subunits of RNA polymerase II were fused to an FRB fragment that can be anchored away from the nucleus by rapamycin addition (Supplementary Fig. 3a,b)^42^. We arrested cells in late G1 phase, by overexpressing the Cdk inhibitor Sic1, when dynamic cohesin turnover on chromosomes is promoted by the cohesin release factor Wapl (Supplementary Fig. 3c)^43^. Following transcription inhibition, loop signals between cohesin binding sites vanished (Fig. 2a,b), confirming a key role of transcription in establishing the loop pattern. When we repeated the experiment in G2/M arrested cells, loop signals substantially weakened following transcription inhibition but remained detectable, consistent with the idea that loops are stabilised by cohesin acetylation during S phase^44,45^.

**Fig. 2.**
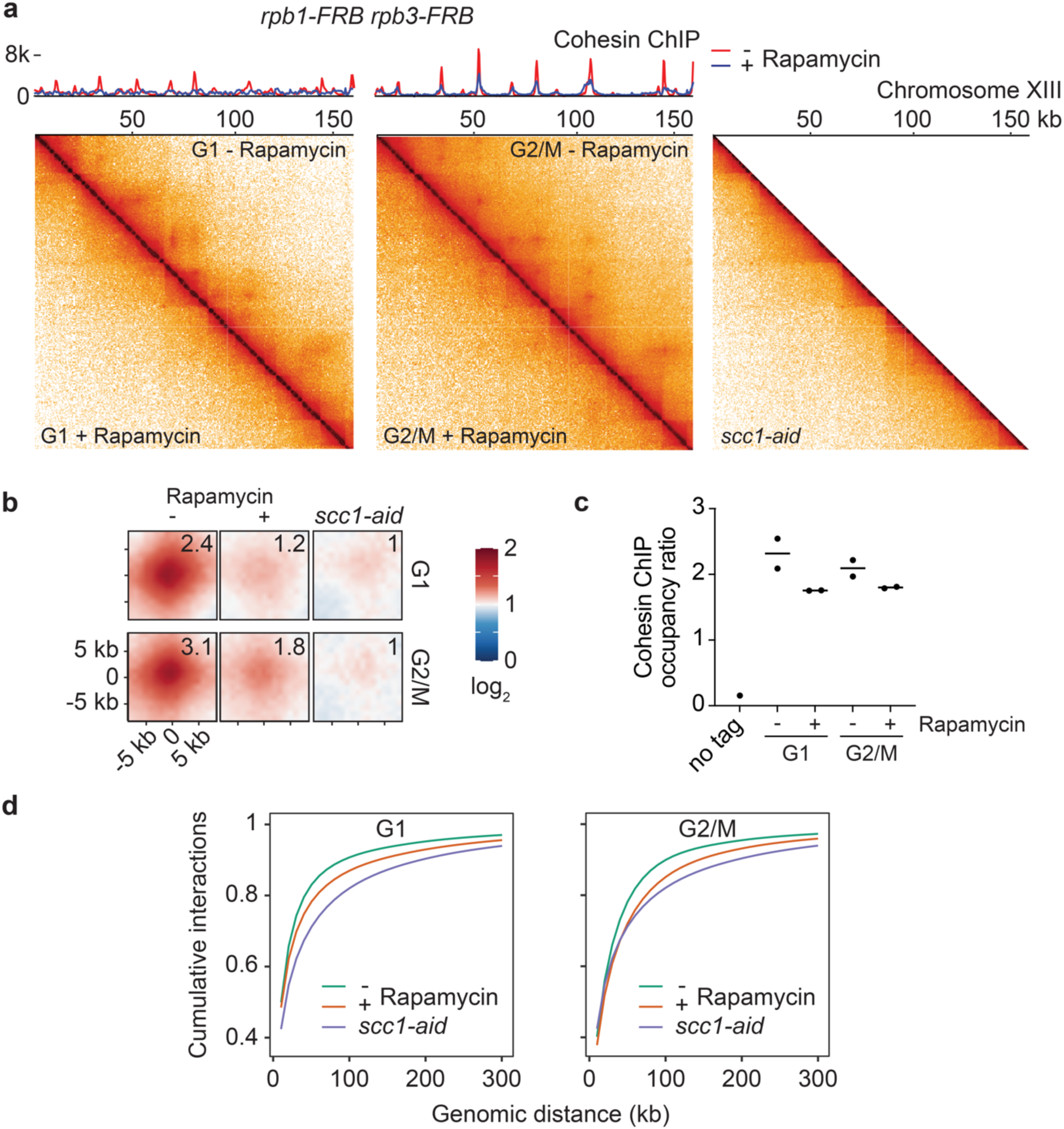
Transcription promotes cohesin loop formation. **a,** 500 bp-resolution merged micro-C contact maps from two independent experiments of cells arrested in late G1 or G2/M in the absence or presence of rapamycin to anchor away RNA Polymerase II. Calibrated cohesin (Smc3-Pk_3_) ChIP-seq traces from the same samples are shown. A merged micro-C contact map of two independent repeats of cohesin-depleted (*scc1-aid*) G2/M cells is shown for comparison. **b,** Aggregate chromatin profiles detected by chromosight and linked to cohesin anchors in the - rapamycin samples (n = 788 G1; n = 1060 G2/M). Mean corner scores are shown relative to those recorded in the *scc1-aid* sample. **c,** Overall cohesin ChIP occupancy in the two repeat experiments, relative to a *C. glabrata* spike-in. **d,** Cumulative interaction counts as a function of genomic distance in the above micro-C experiments.

As a control for the looping pattern, calibrated ChIP-sequencing showed that cohesin remained detectable at most of its usual binding sites, but peaks appeared less sharp, confirming previous observations following chemical transcription inhibition (Fig. 2a)^40^. Overall cohesin levels on chromosomes were only slightly reduced following transcription inhibition (Fig. 2c). Cohesin occupancy at loop anchors was reduced, in both G1 and G2/M cells, however cohesin occupancy declined far less steeply than loop intensities (Supplementary Fig. 3d). These observations open the possibility that transcription promotes chromatin loop formation by a mechanism additional to recruiting and positioning cohesin.

The idea that transcription blocks loop extrusion stems from the observation of longer-ranging intra-chromosome interactions following transcription inhibtion^40,41^ – i.e. cohesins would extrude longer loops when unobstructed by RNA polymerase barriers. However, it remains unknown whether longer-range chromatin contacts following transcription inhibition are indeed cohesin-mediated. To answer this question, we compared chromatin contact distributions following transcription inhibition with those following cohesin depletion. Cumulative interaction plots confirmed a shift toward longer-ranging interactions following transcription inhibition, but also revealed an even more pronounced shift toward longer interactions after cohesin depletion (Fig. 2d). This result can be explained if local, cohesin-mediated contacts are lost following either transcription inhibition or cohesin depletion. As contact frequency distributions reported in Hi-C maps are normalised to unity, loss of local interactions results in a relative, but not necessarily absolute, increase in longer-range interactions. These considerations are consistent with the possibility that transcription cooperates with cohesin during loop formation, but they leave open the question whether RNA polymerases restrict loop growth.

## Transcription promotes, not hinders, loop growth

To investigate whether transcription acts as a loop formation barrier, we utilised a temperature sensitive mutation, *rat1-1*, in the torpedo 5‘-3’ RNA exonuclease that prompts transcription termination at polyadenylation signals^46^. Transcriptional readthrough after *rat1-1* inactivation displaces a subset of cohesin peaks and leads to cohesin accumulation at adjacent peaks^47^. If transcription restricts loop extrusion, pervasive transcription following *rat1-1* inactivation should impose additional constraint and result in smaller loop sizes. Contrary to this prediction, the micro-C pattern in *rat1-1* cells at a restrictive temperature of 37 °C revealed many longer chromatin loops (Fig. 3a). A summary of instances where cohesin peaks are lost due to pervasive transcription is shown in Fig. 3b, exemplifying chromatin loop size increases in all cases. These observations suggest that transcription does not hinder chromatin loop growth but, on the contrary, promotes it.

**Fig. 3.**
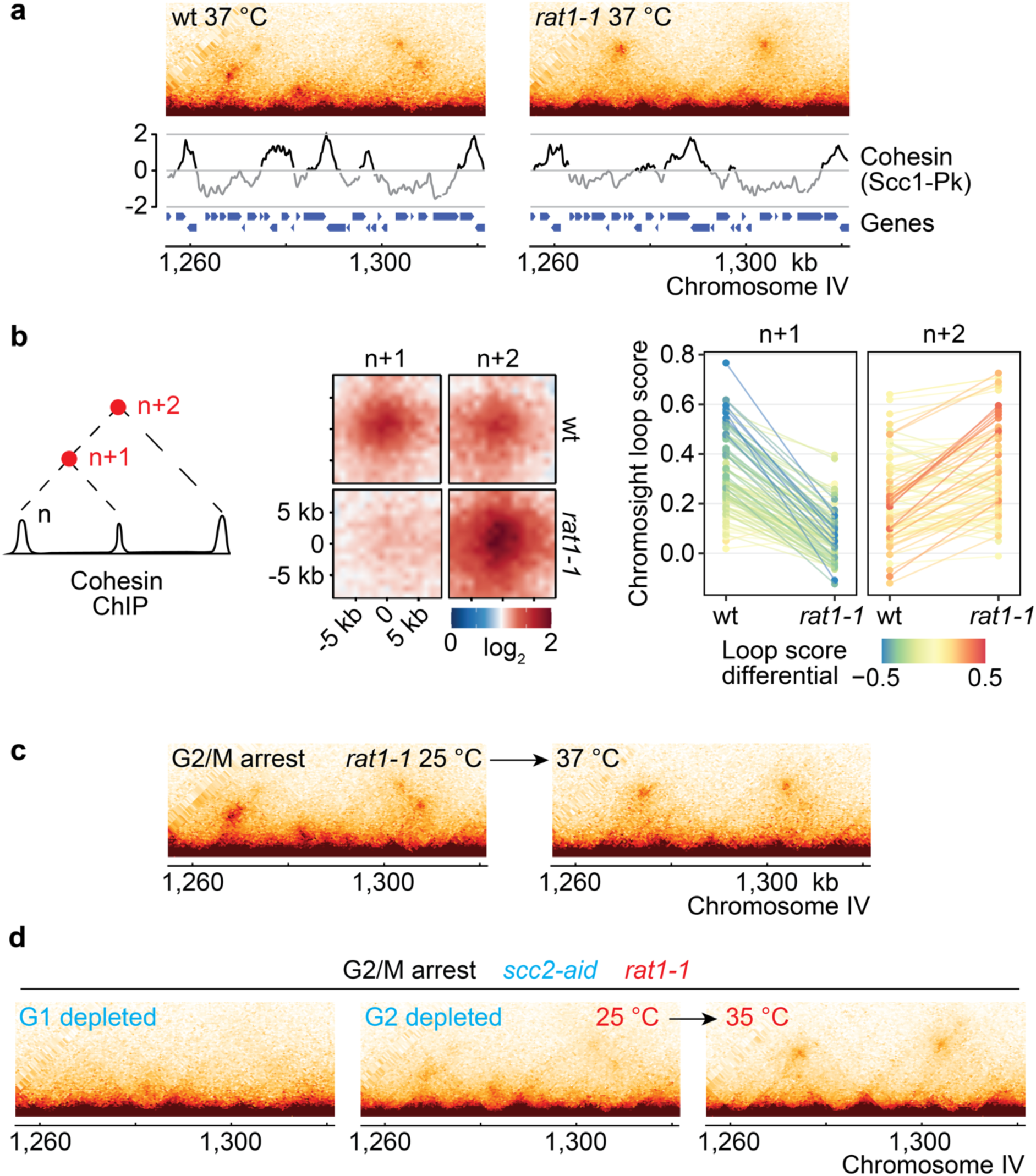
Transcription enlarges cohesin loops. **a,** 500 bp-resolution merged micro-C contact maps from two independent experiments of wt and *rat1-1* cells arrested in G2/M at a restrictive temperature (37 °C) for the *rat1-1* allele. Cohesin (Scc1-Pk_9_) ChIP-microarray traces under the same conditions^47^ are shown. **b,** Scheme for how genomic regions were selected for analysis, aggregate loop profiles, and *rat1-1* dependent change of chromosight loop scores at each position (n=104). **c,** The experiment in panel **a** was repeated, but *rat1-1* cells were arrested in G2/M at a permissive temperature (25 °C, left), before temperature shift to 37 °C (right). **d,** The experiment in panel **c** was repeated with a *rat1-1* strain from which the Scc2 cohesin loader subunit could be depleted. Scc2 was either depleted in G1 before release and arrest in G2/M (left), or following arrest in G2/M. Following G2/M depletion, samples were analysed at 25 °C (middle) and after *rat1-1* inactivation by temperature shift to 35 °C (right).

The above experiment compared steady state loop sizes between wild type and *rat1-1* cells. We next investigated whether pre-existing chromatin loops change their size following *rat1-1* inactivation. We arrested *rat1-1* cells at a permissive temperature of 25 °C in G2/M, when existing chromatin loops have stabilised. *rat1-1* inactivation by temperature shift led to loop expansion, yielding a final loop pattern similar to that seen above (Fig. 3c and Supplementary Fig. 4a). This outcome suggests that transcription enlarges pre-existing chromatin loops.

## Loop growth without loop extrusion

The observation that pervasive transcription expands cohesin-linked chromatin loops allowed us to perform a final test to address how cohesin’s *in vitro* loop extrusion activity relates to *in vivo* loop growth. *In vitro* loop extrusion depends on the cohesin loader complex (Fig. 1b)^12–14^. We therefore repeated the loop expansion experiment, but depleted the Scc2 cohesin loader subunit by promoter shut-off and an auxin-inducible degron (Supplementary Fig. 4b)^48^, before *rat1-1* inactivation by temperature shift. When Scc2 was depleted in the same way before cell cycle entry, no observable loops formed, confirming effective depletion (Fig. 3d). In contrast, when we depleted Scc2 after cohesin-dependent loops had formed, *rat1-1* inactivation again resulted in widespread loop expansion throughout the budding yeast genome (Fig. 3d and Supplementary Fig. 4b). These observations suggest that cohesin’s intrinsic, *in vitro*-observed loop extrusion activity is dispensable for *in vivo* chromatin loop growth. Instead, transcription acts as an extrinsic motor that extends pre-exiting chromatin loops, likely by pushing cohesins that are engaged in loop interactions along transcription units^47,49,50^.

## TAD formation without loop extrusion

In addition to loops, TADs are a prominent cohesin-dependent feature of the mammalian interphase chromatin landscape^6,7^. The loop extrusion hypothesis posits that TADs arise as cohesins bring distal DNA segments into proximity while moving along the chromatin chain^8–11^. On the other hand, oligopaint approaches revealed that TADs form independently of cohesin in individual cells^51^, but that cohesin establishes reproducible boundaries that make TADs visible to population-based techniques. Above, cohesin depletion in G1 arrested budding yeast cells caused chromatin loop loss, while TAD structures appeared to persist (Fig. 2a). TAD persistence following cohesin depletion can be also seen in two previously published studies^37,44^. Other chromatin interactions, possibly including depletion attraction^52^, might suffice to maintain the relatively small budding yeast TADs.

We next investigated cohesin’s contribution to *de novo* TAD formation. We turned to the budding yeast *GAL7*-*GAL10*-*GAL1* locus, encompassing a cluster of three galactose-inducible genes. Cells grown in the presence of glucose, when cluster expression is repressed, show inconspicuous chromatin features at this locus. In contrast, a pronounced TAD boundary is observed in cells grown in galactose-containing medium when cluster expression is switched on (Fig. 4a). Next, we again depleted cohesin and only then added galactose to induce cluster expression (Supplementary Fig. 5). As expected, chromatin loop signals were no longer observed in cohesin’s absence. Yet, a prominent TAD boundary formed at the *GAL7*-*GAL10*-*GAL1* locus following galactose addition (Fig. 4b). The insulating property of a strongly expressed gene cluster, as previously seen in bacteria^53^, appears sufficient to enact a domain boundary. These observations suggest that cohesin, and by inference cohesin-mediated loop extrusion, is not essential for TAD formation in budding yeast.

**Fig. 4.**
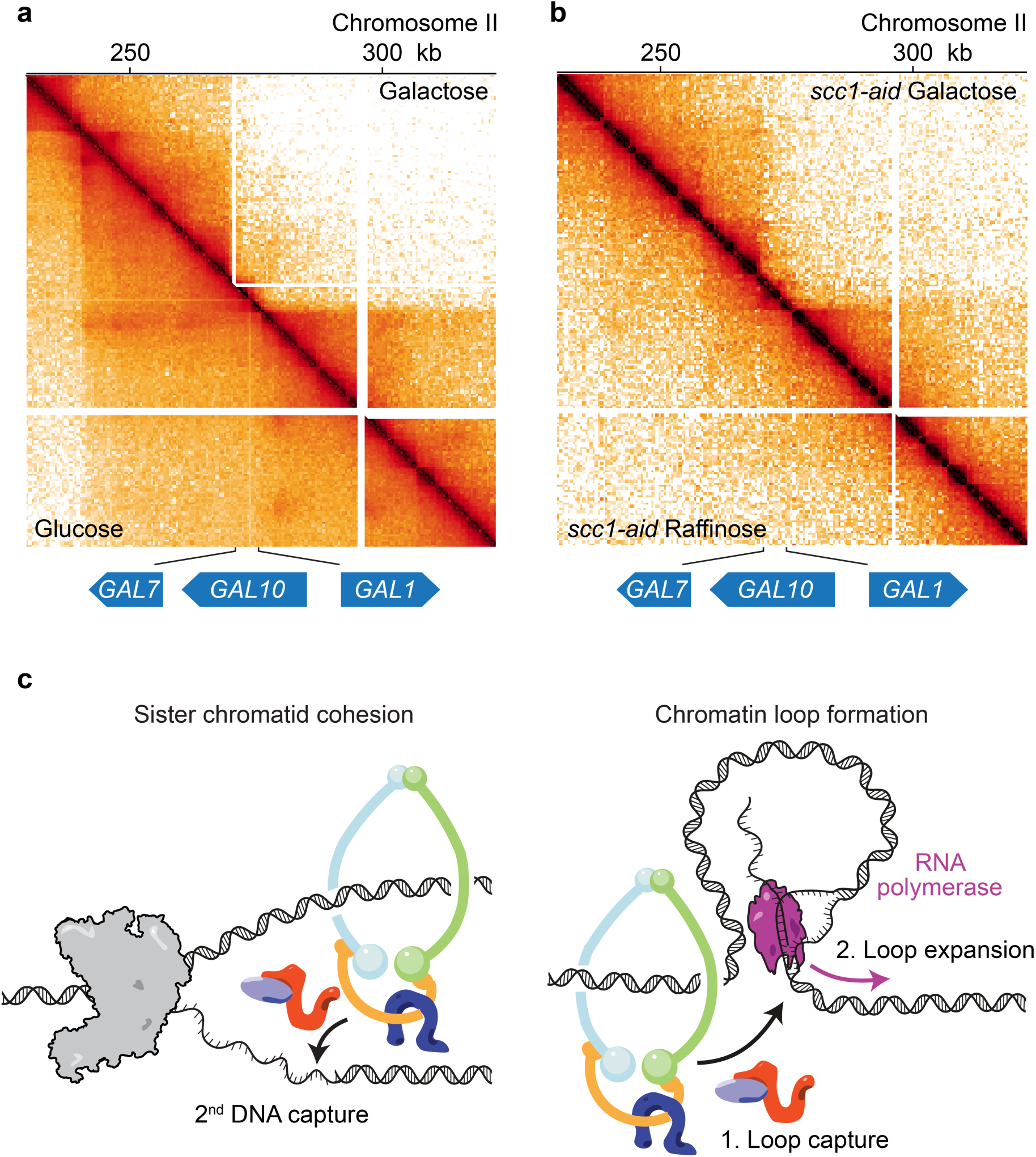
TAD formation without cohesin. **a,** 500 bp-resolution merged micro-C contact maps from two independent experiments surrounding the *GAL7-GAL10-GAL1* locus of cells grown in medium with glucose, or with raffinose + galactose (Galactose) as the carbon source. **b,** as panel **a**, but cells were grown in medium containing raffinose as the carbon source. The cohesin subunit Scc1 was depleted by an auxin-inducible degron. Samples were then analysed in raffinose medium, or following galactose addition. **c,** A model that unifies sister chromatid cohesion and chromatin loop establishment by the cohesin complex.

## Discussion

The loop extrusion hypothesis has shaped current thinking about chromosomal processes. The concept is supported by *in vitro* observations, but had not yet been experimentally tested *in vivo*. We have now studied cohesin variants that lost their *in vitro* loop extrusion ability, and we have removed a key component of the loop extrusion mechanism, the cohesin loader. In both cases, phenomena ascribed to loop extrusion, the formation and growth of chromatin loops, remained unchanged. While *in vitro* loop extrusion constitutes a striking phenomenon, which under certain conditions might arise as by-product of the topological cohesin loading reaction^14,54^, our observations suggest that *in vivo* chromatin loops form by a different mechanism.

Instead of loop extrusion, we find that transcription promotes loop formation and acts as an extrinsic motor that expands chromatin loops. While we performed our study using the simple budding yeast model, evidence for transcription-assisted loop formation is apparent in higher eukaryotes: cohesin promotes sister chromatid cohesion during early vertebrate development, while interphase chromatin structure emerges only once transcription commences during zygotic genome activation^38^. Loops and TADs are lost every time human cohesin dissociates from chromatin during cell divisions, and their re-establishment depends on transcription^39^. Combined single cell Hi-C and RNA-sequencing revealed that most chromatin interaction changes during developmental transitions coincide with, or follow, transcription changes^55^, with the exception of enhancer-promoter interactions that precede transcription changes but are established in a cohesin-independent, stochastic search pattern^20,56,57^. Other studies did not observe a transcription requirement for cohesin chromatin loop formation^58^, though we note the experimental challenge of efficiently shutting down RNA polymerases. Those studies that reported a transcription requirement for loop formation used increased inhibitor concentrations or combined more than one tool to suppress transcription^39,40^.

How might transcription promote chromatin loop formation? Transcription bubbles are DNA structures that bear resemblance to replication forks. The unwound non-template DNA strand, or regions left open by negative supercoiling in the wake of the RNA polymerase, are accessible to ssDNA binding proteins^59^. They might also be accessible for second DNA capture by cohesins that were previously topologically loaded close-by, e.g. at nucleosome-free promoter regions^48,60–62^ (Fig. 4c). While the molecular nature of cohesin’s transcription-dependent loop substrate remains to be further explored, chromatin contact stripes along active mouse genes are suggestive of cohesin-mediated interactions between open promoter elements and the moving transcription machinery^58^. Such stripes are exemplified in nematodes in the form of ‘jets’ that stretch out from cohesin loader binding sites along active genes^63^. If cohesin is stable enough, e.g. following downregulation of its release factor Wapl, this process could extend over long distances, as in the case of locus scanning during VDJ or class switch recombination^64^. Loop establishment by transcription assisted DNA-DNA capture naturally unifies cohesin’s two activities in sister chromatid cohesion establishment and chromatin loop formation.

A mechanism for loop formation in which loop anchors find each other by Brownian diffusion, as proposed here, can explain the simultaneous establishment of both short, as well as of much longer loops, following cohesin depletion and re-addition in mouse cells^6^. This mechanism can also explain generation of both stem-and circle-loops in the fruit fly, the latter of which cannot form by loop extrusion^65^. A transcription-aided loop capture mechanism makes the strong prediction that cohesin is not restricted to intra-chromosome interactions but, at least occasionally, establishes contacts between chromosomes. The chromatin interaction spectrum of fission yeast cohesin, like that of the related condensin complex, indeed encompasses both intra- and inter-chromosome contacts^66,67^. The findings presented in our study motivate a rethinking of how the chromosomal cohesin complex shapes the interphase genome.

## Methods

### Yeast culture

All yeast strains used in this study were of W303 background. *SCC3*^3E^- and *SMC1*^4E^-strains were generated by altering the endogenous gene loci using gene targeting constructs. Successful targeting was confirmed by PCR-based genotyping and DNA sequencing. Cells were grown at 30 °C in YPD medium, except if stated otherwise. Asynchronous mid-log phase cells were diluted to an optical density of OD_600_ = 0.2. Cell synchronisation in G1 was achieved by the addition of 7.5 μg/ml of the mating pheromone α-factor, every hour, for two hours. For late G1 arrest, after cohesin subunit Scc1 expression commences but before the onset of DNA replication^43^, cells were grown in YP medium containing 2% raffinose as the carbon source. Sic1^V5,V33,A76^ expression was induced by 2% galactose addition at the same time as α-factor addition, and the arrest was extended to three hours. Cells were then washed by filtration and released in raffinose and galactose-containing medium without α-factor. G2/M arrest was achieved by releasing cells from α-factor synchronisation into YPD medium containing 6 μg/ml nocodazole. For RNA polymerase II anchor away, 1 μg/ml rapamycin was added for 2 hours before analysis. Scc2 depletion was achieved by addition of 2 mM methionine to shut off *MET3* promoter expression, as well as 1 mM auxin (indole-3-acetic acid). For experiments with the *rat1-1* strain, cells were grown at 25 °C and, as indicated, the temperature was shifted to 37 °C in a water bath, or gradually raised to 35 °C in an air incubator.

### Cell cycle profiling by flow cytometry

Cells were harvested and fixed in 70% ice-cold ethanol for at least 2 hours before pelleting and resuspension in 50 mM Tris-HCl pH 7.5 including 0.1 mg/ml RNase A and incubation overnight at 37 °C. DNA was stained with 25 μg/ml propidium iodide in 200 mM Tris-HCl pH 7.5, 210 mM NaCl, 78 mM MgCl_2_. Cells were sonicated and diluted in 50 mM Tris-HCl pH 7.5. Flow cytometric analyses were performed using an LSRFortessa X-20 flow cytometer (BD Biosciences). 10,000 cells were counted for each sample. Results were visualised using FlowJo.

### Cell viability assays

Asynchronously growing yeast cultures were adjusted to equal optical densities, then 3 μl drops of 10-fold serial dilutions were applied to YPD agar plates, to which supplemental chemicals had been added at the indicated concentrations. Plates were incubated at 30 °C degrees for 2 days.

### Sister chromatid cohesion assay

Strains expressing a tetR-GFP fusion protein and harbouring tetO repeats integrated at the *URA3* locus^16^ were synchronised by α-factor treatment and released for 2 hours into YPD medium containing 6 μg/ml nocodazole. Culture aliquots were harvested and fixed in 70% cold ethanol overnight. Cells were resuspended in PBSA, sonicated, and mounted on 2% agarose patches prepared on glass slides. Z-stacks of 20 images at 0.25-μm intervals were acquired using a DeltaVision Olympus IX70 inverted microscope equipped with a 100× (NA = 1.40) PlanApo objective, deconvolved, and merged using maximum intensity projection. The number of visually discernible GFP dots in each cell were then manually counted. More than one GFP dot was taken to indicate defective sister chromatid cohesion. One hundred cells were counted for each strain, experiments were three times independently repeated.

### Calibrated ChIP-sequencing analysis

50 OD_600_ units of *Saccharomyces cerevisiae* cells were mixed at a 5:1 ratio with *Candida glabrata* cells in which the Smc3 subunit was fused to a Pk epitope tag^68^, and broken in 700 μl Lysis Buffer (50mM HEPES-KOH pH 7.5, 140 mM NaCl, 1 mM EDTA, 1% Triton, 0.1% sodium deoxycholate, supplemented with protease inhibitors) by glass bead rupture. The lysate was retrieved by centrifugation for 1 minute at 2,000 rpm and the supernatant then removed after centrifugation for 10 minutes at maximum speed in a benchtop centrifuge. The pellet was resuspended in 1 ml lysis buffer and sonicated to reach an average DNA fragment size of 200 – 300 bp. Cohesin was precipitated using an α-Pk antibody (clone SV5-Pk1, Bio-Rad) and protein A-coupled Dynabeads for 2 hours. Beads were washed 3 times with Lysis Buffer, 3 times with Lysis Buffer supplemented with 500 mM NaCl and 2 times with Wash Buffer (10 mM Tris-HCl pH 8.0, 250 mM LiCl, 0.5% NP-40, 0.5% sodium deoxycholate). The samples were resuspended in 10 mM Tris-HCl pH 8.0, 1 mM EDTA, 1% SDS (including 3 μg/ml proteinase K and 2 μg/ml RNase A) and de-crosslinked overnight at 65 °C. DNA was purified by phenol-chloroform extraction and the aqueous fraction was further cleaned using DNA Clean & Concentrator-5 with Zymo-Spin Columns (Zymo Research). Libraries were built with the NEBNext Ultra II DNA Library Prep Kit for Illumina (New England Biolabs) following the manufacturer instruction. Sequencing was performed on an Illumina NovaSeq 6000 sequencer.

Sequence data analysis was performed using the nf-core/chipseq pipeline (doi: 10.5281/zenodo.3240506)69. Sequences were aligned to the W303 genome (https://www.ncbi.nlm.nih.gov/nuccore/LYZE00000000)^70^. Calibration was then performed as described^68^. ChIP-seq traces were plotted using PyGenomeTracks (3.8)^71^. Cohesin peaks were called using MACS2 (2.2.9.1)^72^ in paired mode with q < 0.01 as threshold.

### ChIP microarray analysis

ChIP microarray analysis was performed using Affymetrix GeneChip Yeast Genome 2.0 arrays as described^73^. Output files were then transposed from *S. cerevisiae* S288C to W303 genome annotations using a liftOver chain (http://hgdownload.soe.ucsc.edu/admin/exe/).

### Micro-C analysis

Micro-C was performed following published protocols^37,74^ with minor modifications. Yeast cultures were crosslinked with 3% formaldehyde for 15 minutes at 30 °C. The reactions were quenched by the addition of 250 mM glycine at 30 °C for 5 minutes with agitation. Cells were pelleted by centrifugation at 4,000 rpm at 4 °C for 5 minutes and washed twice with water. Cells were then resuspended in Buffer Z (50 mM Tris-HCl pH 7.5, 1 M sorbitol, 10 mM β-mercaptoethanol) and spheroplasted by addition of 250 μg/ml Zymolyase (100T) in a 30 °C incubator at 200 rpm for 40 to 60 minutes. Spheroplasts were washed once by 4 °C PBS and then pelleted at 4,000 rpm at 4 °C for 10 minutes. Chromatin was further crosslinked by suspending pellets in PBS supplemented with 3 mM disuccinimidyl glutarate (ThermoFisher) and incubated at 30 °C for 40 minutes with gentle shaking before quenching by addition of 400 mM glycine for 5 minutes at 30 °C. Cells were pelleted by centrifugation at 4,000 rpm at 4 °C for 10 minutes, washed once with ice-cold PBS and stored at −80 °C. Pellets were then treated as described^37^ until the de-crosslinking step. De-crosslinking solution was added with an equal volume of phenol-chloroform-isoamyl alcohol (25:24:1), vortexed intensively and centrifuged for 15 minutes at room temperature. The aqueous phase was purified using a ZymoClean column (Zymo Research) according to the manufacturer’s instructions. Dinucleosome-sized DNA fragments were purified and excised from a 3% NuSieve GTG agarose gel (Lonza) using the Zymoclean Gel DNA Recovery Kit (Zymo Research). Micro-C libraries were then prepared using the NEBNext Ultra II DNA Library Prep Kit for Illumina (New England Biolabs) as described^74^ and sequenced using an Illumina NovaSeq 6000.

Micro-C datasets were processed through the Distiller pipeline (https://github.com/open2c/distiller-nf, commit 8aa86e) to implement read filtering, alignment, PCR duplicate removal, binning and balancing of replicate sample matrices. Reads were aligned to the W303 genome using bwa (0.17.7) and the resulting maps filtered to remove low-quality alignments (MAPQ<30) and cis alignment pairs within 150 bp. Replicates were analysed independently, and their quality assessed before aggregation into sample-level datasets. Maps were visualised and explored using HiGlass^75^. Loops were called using chromosight (https://github.com/koszullab/chromosight)^76^, as described^77^, using a threshold;: 0.3, and intersected with cohesin ChIP-seq peaks. Aggregate profiles were generated using chromosight, with datasets subsampled to the lowest-depth dataset in the comparison. Cooler files were read into R (4.1.1) using HiCExperiment (1.0.0) (doi:10.18129/B9.bioc.HiCExperiment) and loop files read using InteractionSet (1.22.0)^78^. To compare loop intensities, we applied the HiCExperiment corner score metric that measures the interaction differential between the centre and corner of each loop region.

### Cohesin purification

Cells harbouring Smc1, Smc3, Scc1, and Scc3 expression vectors were grown in YP medium containing 2% raffinose to OD_600_ = 1.0 at 30 °C. 2% galactose was added to the culture to induce protein expression for 2 hours. For purification of Scc3^3E^- and Smc1^4E^-cohesin, both the overexpressed and endogenous copies of these subunits carried the respective mutations (Scc3 K423E, K520E, K669E and Smc1 R53E, R58E, N60E, K63E). Cells were collected by centrifugation, washed once with PBSA and resuspended in an equal volume of Lysis Buffer (50 mM Tris-HCl pH 7.5, 300 mM NaCl, 1 mM MgCl_2_, 10% glycerol, 0.5 mM TCEP, 0.5 mM Pefabloc (Roche) and an additional protease inhibitor cocktail (cOmplete, Roche)). The cell suspension was frozen in liquid nitrogen and broken in a freezer mill. The cell powder was thawed on ice and adjusted with two volumes of Lysis Buffer complemented with 50 U/ml benzonase. The lysates were clarified by centrifugation at 30,000 x g for 15 minutes at 4 °C, then at 142,000 x g for 1 hour. The clarified lysate was passed through a HiTrap NHS-Activated

HP affinity column adsorbed with rabbit IgG according to the manufacturer instructions (Cytiva) and washed with Lysis Buffer, Lysis Buffer including 1 mM ATP, and then Lysis buffer including 0.01% NP-40, before overnight incubation with 10 μg/ml PreScission protease. The eluate was loaded onto a HiTrap Heparin HP column (Cytiva) that was developed with a linear gradient from 300 mM to 1 M NaCl in buffer A (50 mM Tris-HCl pH 7.5, 10% glycerol, 0.5 mM TCEP). The peak fractions were pooled and loaded onto a Superose 6 10/300 GL gel filtration column (Cytiva) that was equilibrated and developed with Buffer A containing 150 mM NaCl. The peak fractions were concentrated by ultrafiltration.

### Loader purification

The cohesin loader Scc2-Scc4 complex was purified as described^30^.

### Electrophoretic gel mobility shift assay

Increasing concentrations of cohesin were incubated for 30 minutes with 16.6 nM (molecules) 505 bp dsDNA (a PCR product from yeast genomic DNA) at 30 °C in 40 mM Tris-HCl pH 7.5, 50 mM NaCl, 2 mM MgCl_2_, 0.5 mM ATP, 0.5 mM TCEP. The reactions were then separated by 0.8% agarose/TAE gel electrophoresis. DNA was detected by staining with SYBR Gold for one hour, followed by 15 minutes destaining in TAE before imaging with a Gel Doc XR+ (Bio-Rad) gel documentation system.

### *In vitro* cohesin loading assay

In a reaction volume of 15 μl, 30 nM cohesin, 60 nM Scc2–Scc4 cohesin loader and 3.3 nM (molecules) pBluescript II KS(+) DNA were mixed in 35 mM Tris-HCl pH 7.0, 20 mM NaCl, 0.5 mM MgCl_2_, 13.3% glycerol, 0.5 mM ATP, 0.003% Tween, and 1 mM TCEP. The reactions were incubated at 30 °C for one hour. Reactions were stopped by the addition of 500 μl of IP Buffer 1 (35 mM Tris-HCl pH 7.5, 100 mM NaCl, 10 mM EDTA, 5% glycerol, 0.35% Triton X-100). 4 μg of α-Pk antibody (clone SV5-Pk1, Bio-Rad) were added and incubated on a wheel at 4 °C for 2 hours, before the addition of 40 mg/ml protein A-coupled Dynabeads followed by 30 minutes additional incubation. The beads were washed three times with IP Buffer 1, twice with IP Buffer 2 (35 mM Tris-HCl pH 7.5, 300 mM NaCl, 10 mM EDTA, 5% glycerol, 0.35% Triton X-100) and once with (35 mM Tris-HCl pH 7.5, 100 mM NaCl, 0.1% Triton X-100).

The beads were suspended in 12 μl of elution buffer (10 mM Tris-HCl pH 7.5, 1 mM EDTA, 50 mM NaCl, 0.75% SDS, 1 mg/ml protease K) and incubated at 37 °C for 30 minutes. The recovered DNA was analysed by 0.8% agarose/TAE gel electrophoresis. The gel was stained with SYBR Gold, as above. Gel images were captured, and band intensities quantified using Fiji (2.3.0/1.53q).

### DNA loop extrusion assays

Microfluidic flow cells were prepared as previously described^14^. Flow cells were incubated with 1 μl of α-digoxigenin antibody (Roche, 150 U) diluted in 30 μl TB buffer (40 mM Tris-HCl pH 7.5, 50 mM NaCl) for 10 minutes and washed with 400 μl TB buffer. The surface of the flow cell was further passivated by incubating with 50 μl of Pluronic F127 (Sigma-Aldrich, 1% solution in TB buffer) for 10 minutes followed by washing with 400 μl TB and incubating with 40 μl of β-Casein (Sigma-Aldrich, 10 mg/ml in TB buffer) for 30 minutes. Subsequently the flow cell was washed 4 times with 400 μl TB buffer. 40 μl of 5 pM λ-phage DNA (New England Biolabs), digoxigenin-labelled at both ends^14^, was introduced into the flow cell in TB buffer at a flow rate of 4 μl/min using a syringe pump (Harvard Apparatus, Pico Plus Elite 11). The flow cell was then washed again with 40 μl TB buffer at a flow rate of 4 μl/min. Prior to imaging, the flow cell was equilibrated with 50 μl buffer R (40 mM Tris-HCl pH 7.5, 50 mM NaCl, 2 mM MgCl_2_, 5 mM ATP, 10 mM DTT, 200 nM SYTOX Orange, 0.2 mg/ml glucose oxidase, 35 μg/ml catalase, 4.5 mg/ml dextrose and 1 mg/ml β-casein) at 15 μl/min. Cohesin tetramer complex was pre-mixed with its loader at an equimolar ratio at 500 nM concentration in buffer R on ice. DNA loop extrusion was initiated by flowing in the cohesin-loader mixture at 5 nM concentration in buffer R at a flow rate of 6 μl/min. DNA molecules stained with SYTOX Orange were imaged using a custom-built HILO microscopy setup utilising a 561 nm laser and a Nikon SR HP Apo TIRF 100x/1.49 oil immersion objective by taking snapshots with 100 ms exposure every second for 8 minutes. To determine the efficiency of DNA loop extrusion in the absence of the flow, the flow was stopped, and the flow cell was incubated with cohesin, loader and ATP for 8 minutes. Flow was then resumed, and snapshots taken every second for 1 minute. Images were collected with an Andor Sona sCMOS camera, saved as uncompressed TIFF files and further processed using Fiji. Experiments were performed at room temperature.

### Analysis of DNA loop extrusion experiments

To determine the efficiency of DNA loop extrusion, the number of DNA molecules containing loops was divided by the total number of double-tethered DNA molecules. Single-tethered DNA molecules were excluded from the analysis. Loop extrusion rates were extracted as previously described^14^. The length of DNA molecules stretched with a constant flow was manually measured before loop extrusion, averaged over a 5 second interval, and normalised to 48.5 kb (the length of a λ-phage DNA). Subsequently, the length of DNA outside the loop was measured, frame by frame, and subtracted from the initial DNA length to calculate the loop size. The loop extrusion rate was calculated as the slope of a linear fit to the measurements.

## Acknowledgements

We would like to thank J. Campbell, D. Lang and R. Mitter for bioinformatics support, D. Jackson and the Crick Advanced Sequencing Facility for their assistance, C. Bouchoux for reagents, A. Gammie for the W303 genome file, D. Bentley, L.-Y. Chu, S. Glaser, Y. Kakui and J. Svejstrup for scientific input, K. Dubrana, S. Marcand and the past and present laboratory members for discussions and critical reading of the manuscript. This work was supported by the Wellcome Trust (220244/Z/20/Z to F.U.) and The Francis Crick Institute, which receives its core funding from Cancer Research UK, the UK Medical Research Council, and the Wellcome Trust (cc2125 to M.M.; cc2137 to F.U.).

## Author contributions

T.M.G. and F.U. conceived the study, T.M.G. performed most experiments, C.B. performed the bioinformatics analyses, G.P. and M.M. contributed the *in vitro* loop extrusion experiments. T.M.G. and F.U. wrote the manuscript with input from all co-authors.

## Competing interests

The authors declare no competing interests.

---

**Supplementary Fig. 1.**
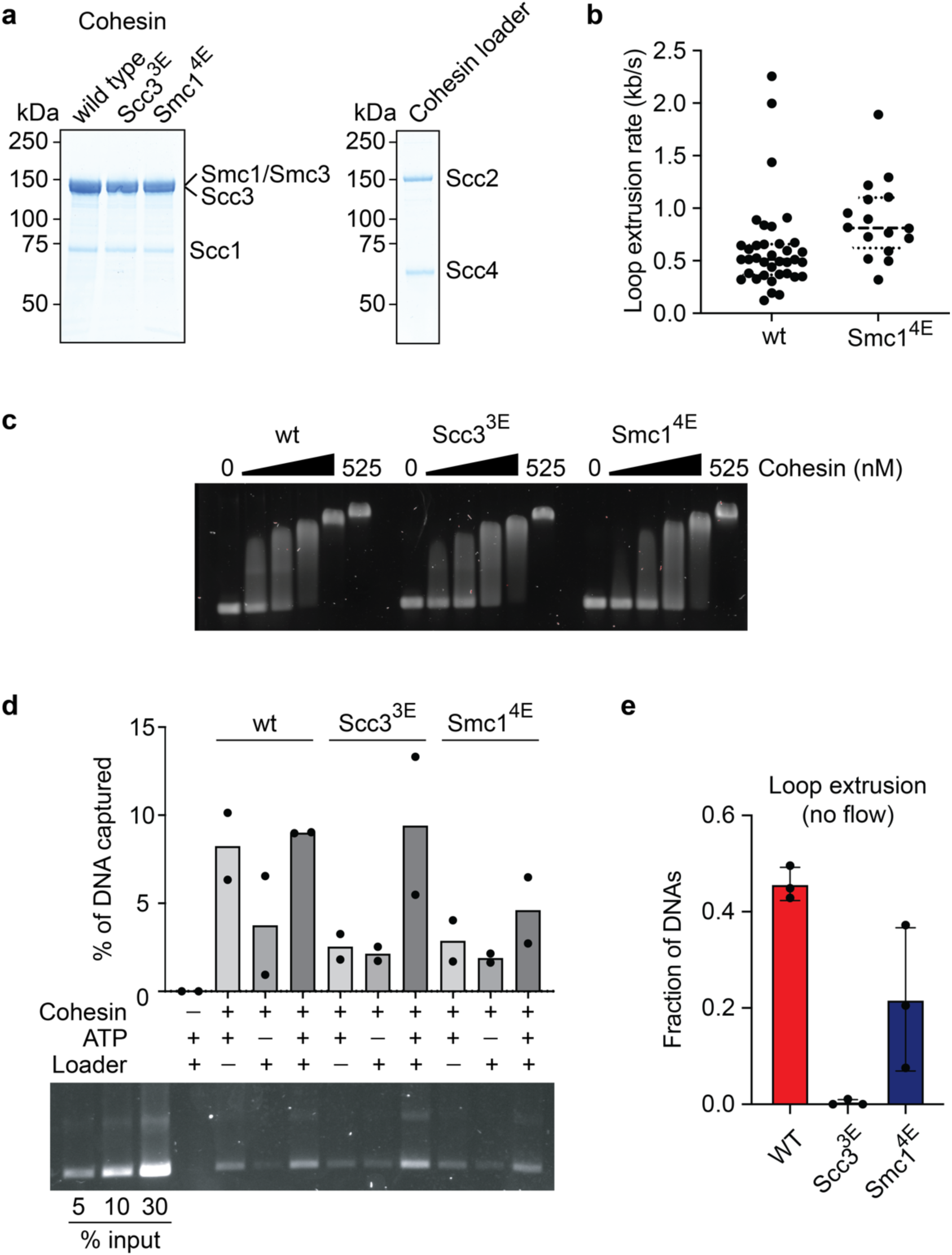
Characterisation of Scc3^3E^ and Smc1^4E^ loop extrusion defective cohesin complexes. **a,** Purified wild type (wt), Scc3^3E^-, and Smc1^4E^-cohesin and cohesin loader were analysed by SDS-PAGE followed by Coomassie Blue staining. **b,** Loop extrusion rates, measured as described^14^, of wt and Smc1^4E^-cohesin, in the presence of loader and ATP (n_wt_ = 37, n_Smc1_^4E^ = 16). Comparable loop extrusion rate distributions suggest that Smc1^4E^-cohesin is defective in loop initiation but less so in loop extension. Dashed and dotted lines represent the median and quartile ranges, respectively. **c,** DNA affinity of wt, Scc3^3E^- and Smc1^4E^-cohesin as measured by an electrophoretic mobility shift assay. **d,** Assay to measure topological (high-salt resistant) loading of wt, Scc3^3E^- and Smc1^4E^-cohesin onto DNA^30^, in the presence of the indicated components. An example agarose gel of the recovered DNA is shown, as well as quantification of the individual results from two independent repeat experiments. Bars show the means. **e,** Loop extrusion assay as in Fig. 1c, but the flow cell was incubated with wt, Scc3^3E^- or Smc1^4E^-cohesin, loader and ATP in the absence of flow, before flow was applied to visualise loops. The fractions of DNA with loops were counted in three independent repeat experiments. Individual data points are shown, bars represent the mean and error bars the standard deviation (n_wt_ = 224, n_Scc3_^3E^ = 269, n_Smc1_^4E^ = 633).

**Supplementary Fig. 2.**
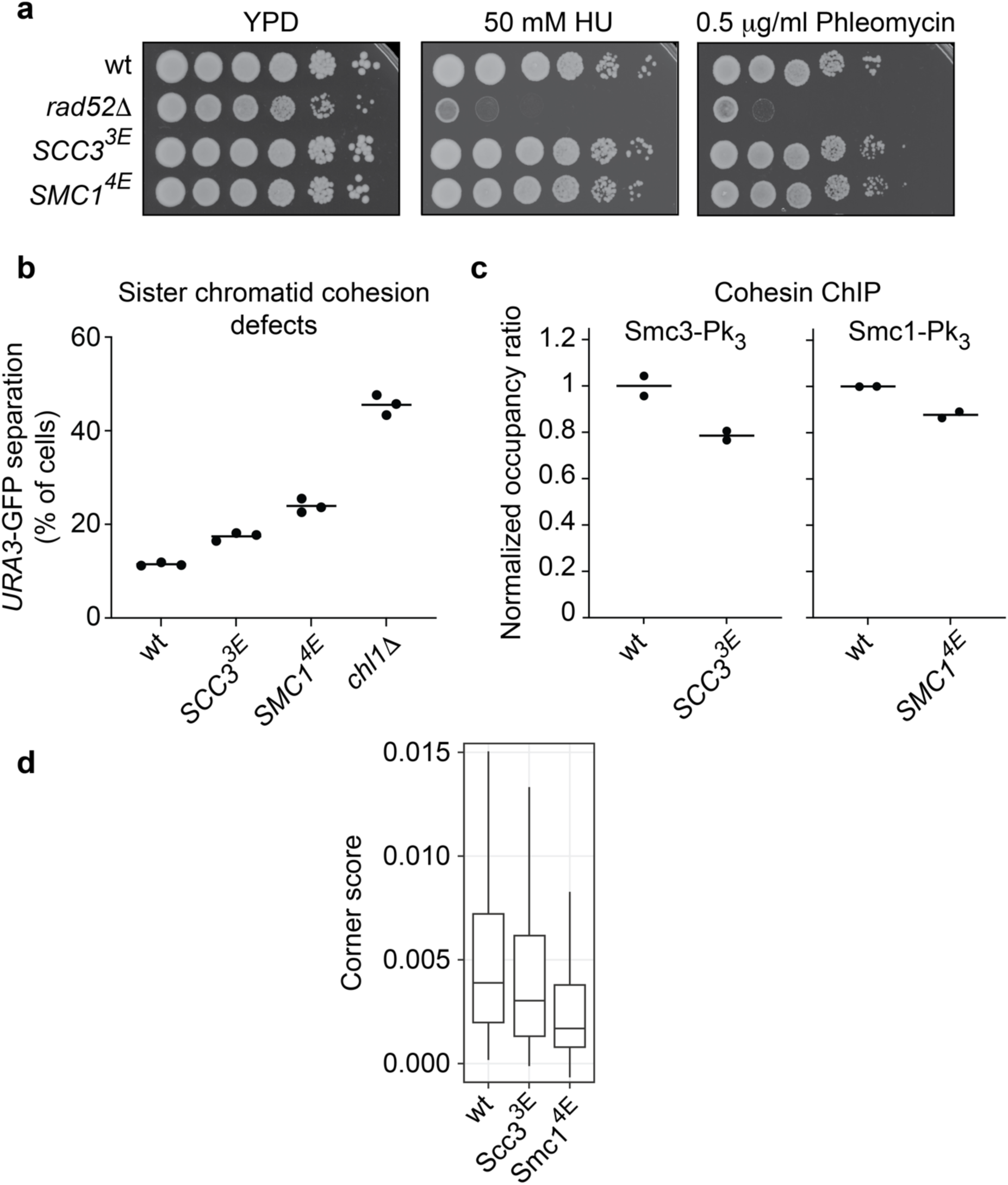
Characterisation of life without loop extrusion. **a,** 10-fold serial dilutions of cultures of the indicated genotypes were plated onto YPD agar plates containing the indicated compounds and grown at 30 °C for 2 days. A wt and a DNA repair deficient (*rad52Δ*) strain were included as controls. **b,** Sister chromatid cohesion in G2/M arrested cells was monitored at the GFP-marked *URA3* locus^16^. The percentage of cells (n = 100) with two separated GFP dots were recorded in three independent repeat experiments. The means are represented by horizontal bars. A wt and a cohesion establishment defective (*chl111*)^80^ strain served as controls. **c,** Overall cohesin ChIP enrichment ratios of wt, compared to Scc3^3E^- and Smc1^4E^-cohesin, relative to a *C. glabrata* spike-in. Cohesin ChIP used Smc3-Pk_3_ in the Scc3^3E^ strain, or Smc1^4E^-Pk_3_, normalised against Smc3-Pk_3_ and Smc1-Pk_3_ wt control strains. **d,** Corner score distributions of loops identified in the wt micro-C contact map, sampled in the Scc3^3E^- and Smc1^4E^-maps.

**Supplementary Fig. 3.**
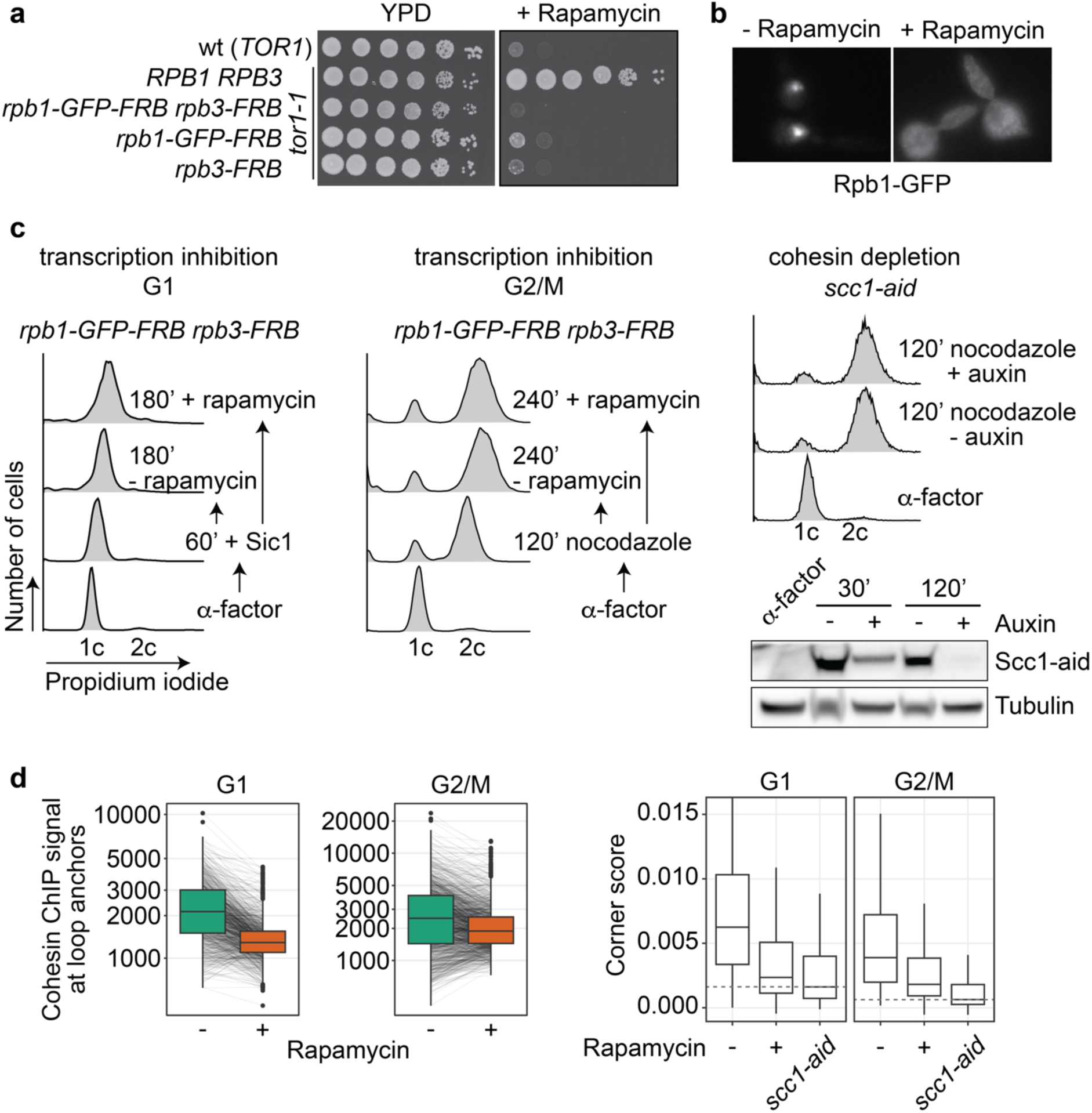
Transcription inhibition and its effect on cohesin-mediated chromatin loops. **a.** 10-fold serial dilutions of cultures of the indicated genotypes were plated onto YPD agar plates, with or without 2 μg/ml added rapamycin, and grown at 30 °C for 2 days. A strain in which both Rpb1 and Rpb3 subunits of RNA polymerase II were fused to FRB showed a tighter response to rapamycin, as compared to strains with either one of the fusions. **b.** An example of Rpb1-GFP-FRB relocation from the nucleus to the cytoplasm after one hour 2 μg/ml rapamycin treatment. **c.** FACS analysis of DNA content of the cells in the experiment shown in Fig. 2, as well as an experimental outline. Western blot analysis confirmed Scc1-aid depletion by its auxin-inducible degron, at 30 minutes and 120 minutes (the time of cell harvest) after release from α-factor synchronisation. **d,** Cohesin ChIP signal intensity distributions at loop anchors (normalised mean reads), in the absence or presence of rapamycin, in both the G1 and G2/M synchronised cultures. Grey lines connect individual ChIP signal intensities under the two conditions. Corner score distributions of the corresponding loops, before and after transcription inhibition, as well as of the same loop positions sampled following Scc1 depletion, are shown alongside. Baseline corner scores in the absence of cohesin are indicated by dashed lines.

**Supplementary Fig. 4.**
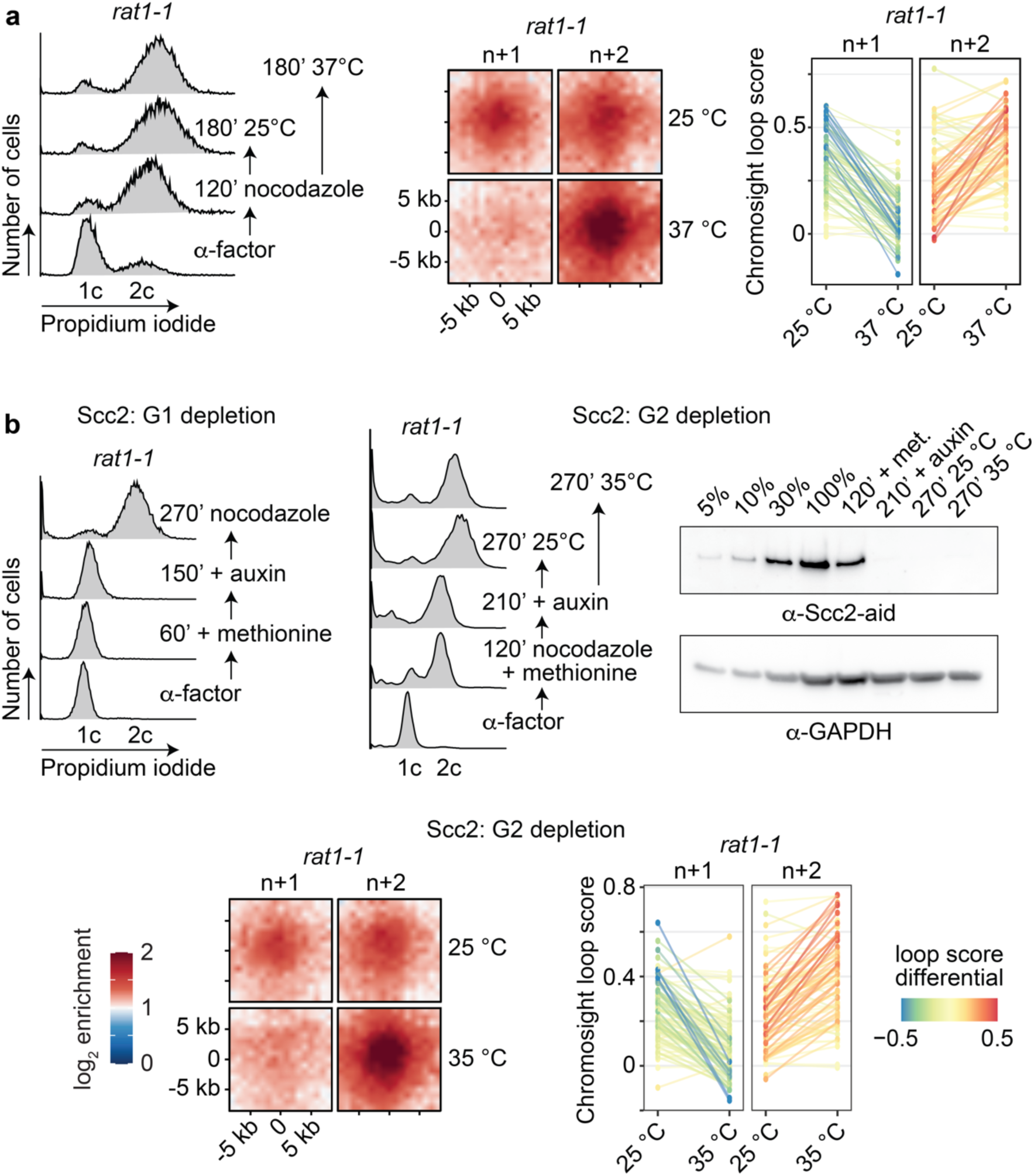
Transcription expands cohesin-mediated chromatin loops. **a.** FACS analysis of DNA content of the cells in the experiment shown in Fig. 3c, together with an experimental outline. Aggregate chromatin profiles of loops (n = 91), identified as in Fig. 3b, and a graph depicting the *rat1-1* dependent loop score changes. **b.** FACS analyses of DNA content of the cells in the experiment shown in Fig. 3d, together with experimental outlines. Western blot analysis confirmed Scc2 depletion by *MET3* promoter repression and an auxin-inducible degron. Serial dilutions of a sample before methionine and auxin addition were loaded, as well as samples at the indicated times in the experiment. Scc2 was detected using an aid-tag antibody. GAPDH served as a loading control. Aggregate loop profiles (n = 97) and a graph depicting the *rat1-1* dependent loop score changes are shown.

**Supplementary Fig. 5.**
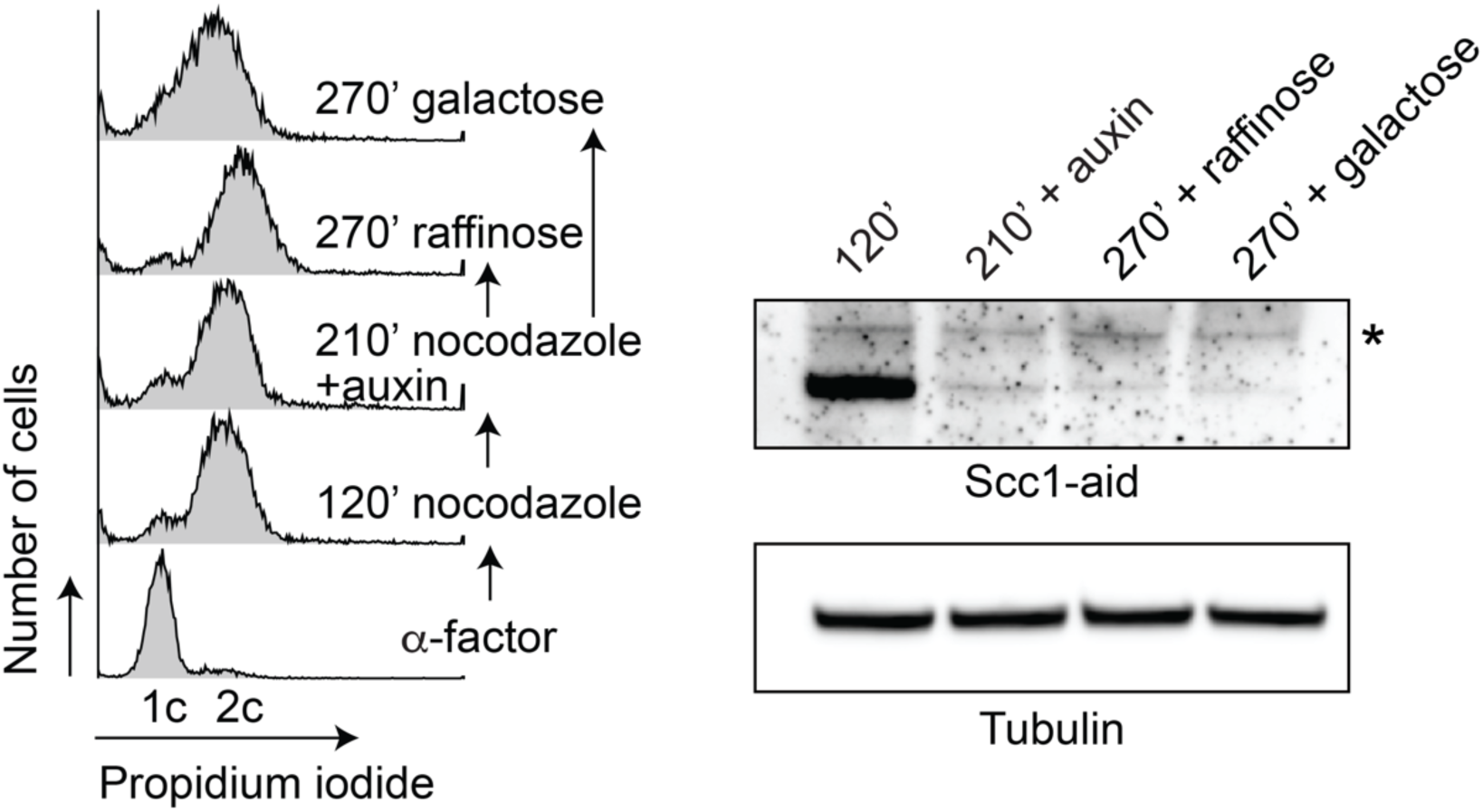
TAD formation without cohesin. FACS analysis of DNA content of the cells in the experiment shown in Fig. 4b, as well as an experimental outline. Western blot analysis confirmed Scc1-aid depletion by its auxin-inducible degron, following auxin addition to a G2/M arrested culture. The asterisk marks a background band recognised by the α-aid-tag antibody. Tubulin served as a loading control.

## Notes

### Competing Interest Statement

The authors have declared no competing interest.

